# *In Vivo* Efficacy and PK/PD Modeling of KBP-7072, An Aminomethylcycline Antibiotic, in Neutropenic Pneumonia and Thigh Infection Models

**DOI:** 10.1101/2020.03.18.998112

**Authors:** Xiaojuan Tan, Min Zhang, Qingmei Liu, Ping Wang, Tian Zhou, Vincent Benn, Fred Yang

## Abstract

KBP-7072 is a novel aminomethylcycline with broad-spectrum activity against Gram-positive and Gram-negative multidrug resistant bacterial isolates and strains. *A*ntibacterial activity and the PK/PD relationship were assessed using *in vivo* infection models. Six to 8-week-old female CD-1 mice were randomized to oral KBP-7072, minocycline and vehicle in a *Klebsiella pneumoniae* murine model, and KBP-7072, linezolid and vehicle for a *Streptococcus pneumoniae* murine model. Each animal was inoculated with *K. pneumoniae* or *S. pneumoniae* placed on the tip of the nares. KBP-7072 and antibiotics were started 3 hours post inoculation and continued for 3 days for *K. pneumoniae*, and were started 18 hours post inoculation and continued for 3 days for *S. pneumoniae*. Animals were euthanized at 0 (control group), 24, 48 or 72 hours post final dose. *In vivo* efficacy and PK/PD parameters were determined in *Staphylococcus aureus* isolate (6424MRSA-363), *K. pneumoniae* isolate (6680kpn-522), and *E. coli* isolate (6691eco-558) murine thigh infection models. *In vivo* efficacy and PK/PD parameters (*f*AUC/MIC, *f*C_max_/MIC and %T>MIC_free_) were calculated. Respiratory infection occurred in all inoculated mice. KBP-7072 produced a significant (*p*<0.05 to <0.001) dose-dependent decrease in colony forming units (CFUs) at all doses and a dose-dependent increase in survival rate (*p*<0.001 vs. vehicle). The median survival in all KBP-7072-treated groups was significantly greater vs. comparators (*p*<0.001). These results demonstrate potent *in vivo* efficacy for KBP-7072 and determined that the AUC/MIC parameter was optimal for assessing bacteriostatic and bactericidal effects of KBP-7072.

## INTRODUCTION

Community-acquired pneumonia (CAP) is the most common infectious diseases leading to hospitalization and mortality among all age groups, but in particular, the young and the elderly.^1–3^ The economic burden of CAP is immense and has been estimated to already exceed $17 billion in the U.S. and $10 billion in Europe.^2,3^ Increasing rates of resistance in recent years among *Streptococcus pneumoniae*, *Haemophilus influenzae*, and other pathogens that are common etiologic agents for CAP impose additional challenges for providing effective treatment.^4–6^ Increased rates of resistance to macrolides^7^ and beta-lactams^8^ have been reported among *S. pneumonia* isolates in the SENTRY program. Others have reported increased rates of bacterial resistance with beta-lactams, fluoroquinolones, macrolides, and earlier generation tetracyclines.^9^ In addition, fluoroquinolone use has been associated with increased risks for tendinitis and tendon rupture, neurological complications, and hypoglycemia that could limit their use for treatment of common infections.^10^ Increasing rates of resistance and potential side effects with some antibiotics, together with the substantial morbidity and mortality associated with CAP emphasize the need for new drugs to add to the treatment armamentarium.

KBP-7072 is a novel, semi-synthetic, aminomethylcycline antibiotic, which inhibits the normal function of the bacterial ribosome. KBP-7072 exhibits a broad spectrum of *in vitro* antibacterial activity against Gram-positive and Gram-negative bacteria including many multidrug resistant pathogens. Notably, KBP-7072 is active against many of the bacteria causing CAP, including *S. pneumoniae*, penicillin-resistant *S. pneumoniae* (PRSP), *H. influenzae*, *Staphylococcus aureus*, methicillin-resistant *S. aureus* (MRSA), vancomycin-resistant enterococcus (VRE), *Enterobacteriaceae* spp, *Acinetobacter* spp, *Pseudomonas* spp. as well as atypical pathogens including *Mycoplasma pneumoniae, Legionella pneumophilia, and Chlamydia pneumoniae*.^11–12^ Results in healthy volunteers administered single and multiple ascending doses demonstrate that KBP-7072 may be administered once daily as an oral and intravenous (IV) formulation.^13,14^

We report results investigating the *in vivo* bactericidal activity of KBP-7072 against *Klebsiella pneumoniae* and *S. pneumoniae* in a murine pneumonia model and the *in vivo* activity and the pharmacokinetic/pharmacodynamic (PK/PD) relationship of KBP-7072 in a mouse thigh infection model.

## RESULTS

### Pneumonia Infection Models

Respiratory infection with *K. pneumoniae* occurred in all inoculated mice. Treatment with KBP-7072 resulted in a significant dose-dependent decrease in CFUs at all doses and most time points tested. At 72 hours post dose, KBP-7072 150, 300, and 600 mg/kg qd, or 75 and 150 mg/kg bid, resulted in significant (*p*<0.05 to <0.001) reductions in bacterial growth (CFUs) vs. vehicle (**Table 1 and Figure 1**). Minocycline qd or bid resulted in no significant reduction of bacterial growth. Treatment with KBP-7072 resulted in a dose-dependent prolonged survival rate (**Figure 2**). At doses of 150 and 300 mg/kg, KBP-7072 resulted in a median survival of 3 days compared to a median survival of 2 days in vehicle-treated animals (p<0.01 to *p*<0.001). Minocycline 300 mg/kg resulted in no increase in median survival, and all animals in vehicle and minocycline groups died by Day 4.

**Table 1.** Mean ± standard deviation CFU at 24, 48, and 72 hours post dose of drug or vehicle in the *K. pneumoniae* and *S. pneumoniae* models.

| ***K. pneumoniae* Model** |  |  |  |
| --- | --- | --- | --- |
| Treatment Group | 24 hours | 48 hours | 72 hours |
| KBP-7072 150 mg/kg qd | 8.60 ± 0.56 | 9.14 ± 1.02 | 8.00 ± 1.72 |
| KBP-7072 300 mg/kg qd | 8.69 ± 0.51 | 8.44 ± 1.11 | 7.64 ± 1.31 |
| KBP-7072 600 mg/kg qd | 7.78 ± 0.50 | 6.38 ± 1.22 | 5.18 ± 0.66 |
| KBP-7072 75 mg/kg BID | 8.62 ± 0.39 | 8.14 ± 1.16 | 7.42 ± 1.25 |
| KBP-7072 150 mg/kg BID | 7.61 ± 0.55 | 7.11 ± 0.64 | 5.12 ± 0.73 |
| Minocycline 300 mg/kg qd | 8.99 ± 0.13 | 9.26 ± 0.55 | 9.43 ± 0.48 |
| Minocycline 150 mg/kg BID | 8.73 ± 0.35 | 8.82 ± 0.51 | 9.41 ± 0.43 |
| Vehicle | 9.46 ± 0.24 | 10.13 ± 0.09 | 9.79 ± 0.48 |
| ***S. pneumoniae* Model** |  |  |  |
| Treatment Group | 24 hours | 48 hours | 72 hours |
| KBP-7072 5 mg/kg qd | 6.16 ± 0.63 | 4.35 ± 1.37 | 3.49 ± 1.90 |
| KBP-7072 15 mg/kg qd | 5.37 ± 0.72 | 3.03 ± 0.88 | 0.40 ± 0.98 |
| KBP-7072 45 mg/kg qd | 4.66 ± 0.28 | 1.52 ± 0.78 | 0 ± 0 |
| KBP-7072 2.5 mg/kg BID | 5.81 ± 0.98 | 2.17 ± 1.10 | 0 ± 0 |
| KBP-7072 7.5 mg/kg BID | 4.71 ± 1.19 | 1.00 ± 1.55 | 0 ± 0 |
| Linezolid 7.5 mg/kg BID | 7.32 ± 0.43 | 8.70 ± 0.47 | 9.12 ± 0.17 |
| Linezolid 15 mg/kg BID | 7.03 ± 1.21 | 7.00 ± 0.72 | 5.94 ± 3.02 |
| Vehicle | 8.19 ± 0.49 | 8.38 ± 1.16 | 8.99 ± 0.50 |

**Figure 1.**
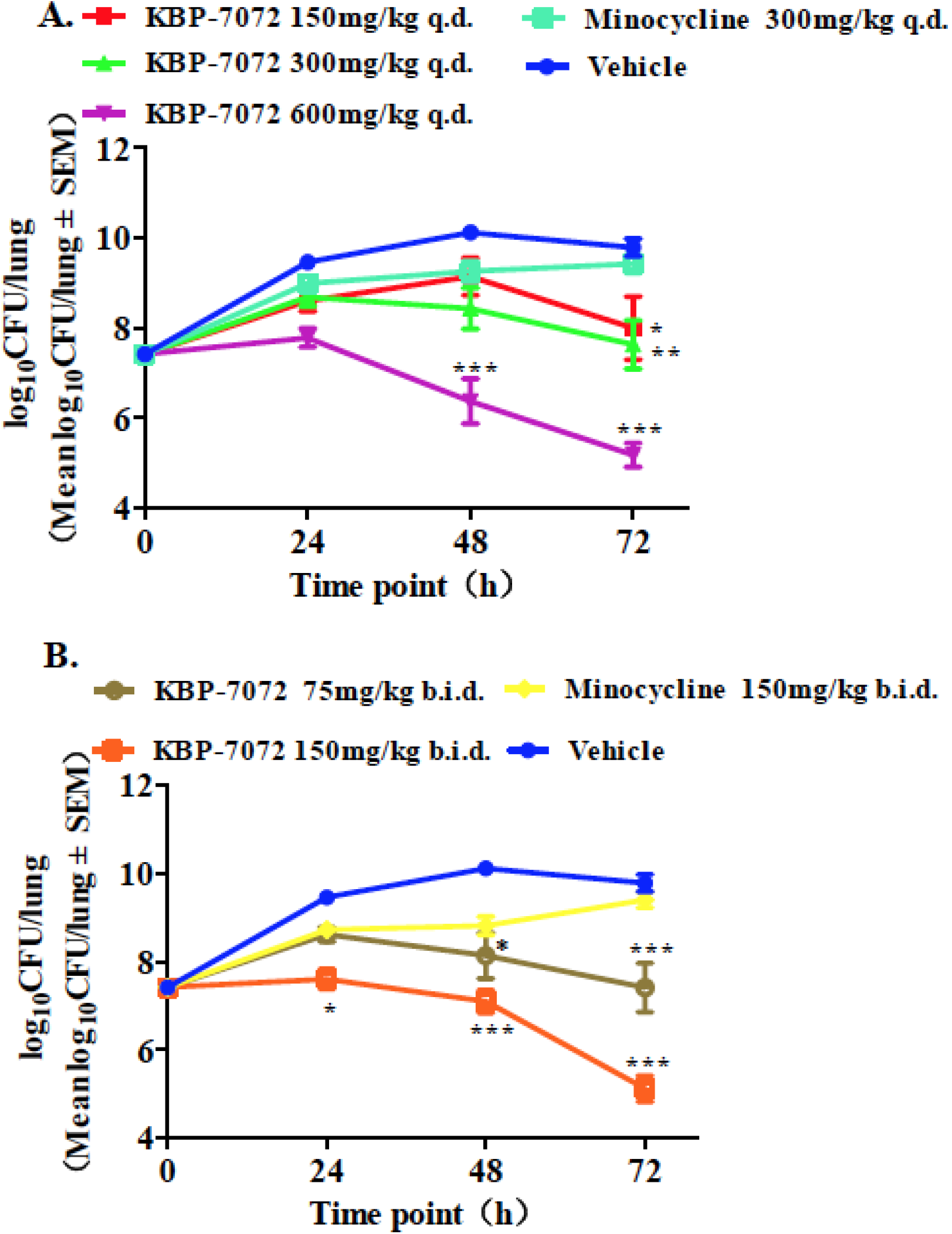
Efficacy of KBP-7072 in mouse pneumonia model infected by intranasal inoculation of *Klebsiella pneumoniae*. A. Colony forming unit in animals treated by KBP-7072 dosed once daily. B. Colony forming unit in animals treated by KBP-7072 dosed bid. (* p<0.05; ** p< 0.01; *** p<0.001).

**Figure 2.**
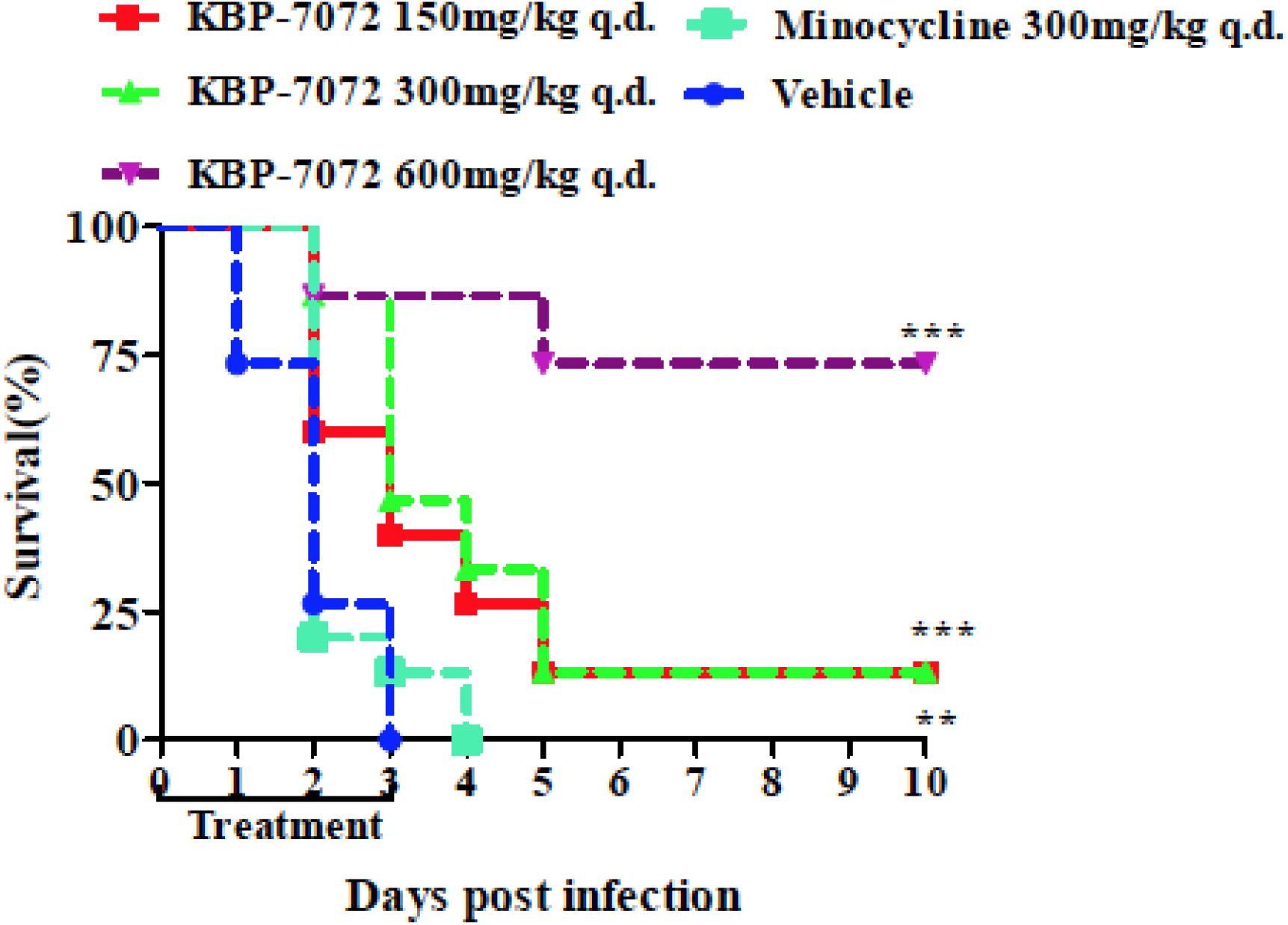
Survival of experimental animals. Log Rank Test was used to analyze survival rate. (* p<0.05; ** p<0.01; *** p<0.001).

In the *S. pneumoniae* model at 72 hours post dose, KBP-7072 45 mg/kg qd, or 2.5 and 7.5 mg/kg bid resulted in no detectable bacteria in the lung and marked lower mean CFUs at 5 and 15 mg/kg doses (**Table 1 and Figure 3**). No significant reduction of bacterial growth occurred with linezolid 7.5 mg/kg bid, but linezolid 15 mg/kg bid resulted in a significant reduction of the number of bacteria in lung vs. vehicle (*p*<0.001). KBP-7072 doses of 5 and 15 mg/kg, resulted in a significant, dose-dependent increases (*p*<0.001) in median survival, and linezolid 7.5 mg/kg resulted in a significant increase (*p*<0.001) in survival compared to vehicle (**Figure 4**).

**Figure 3.**
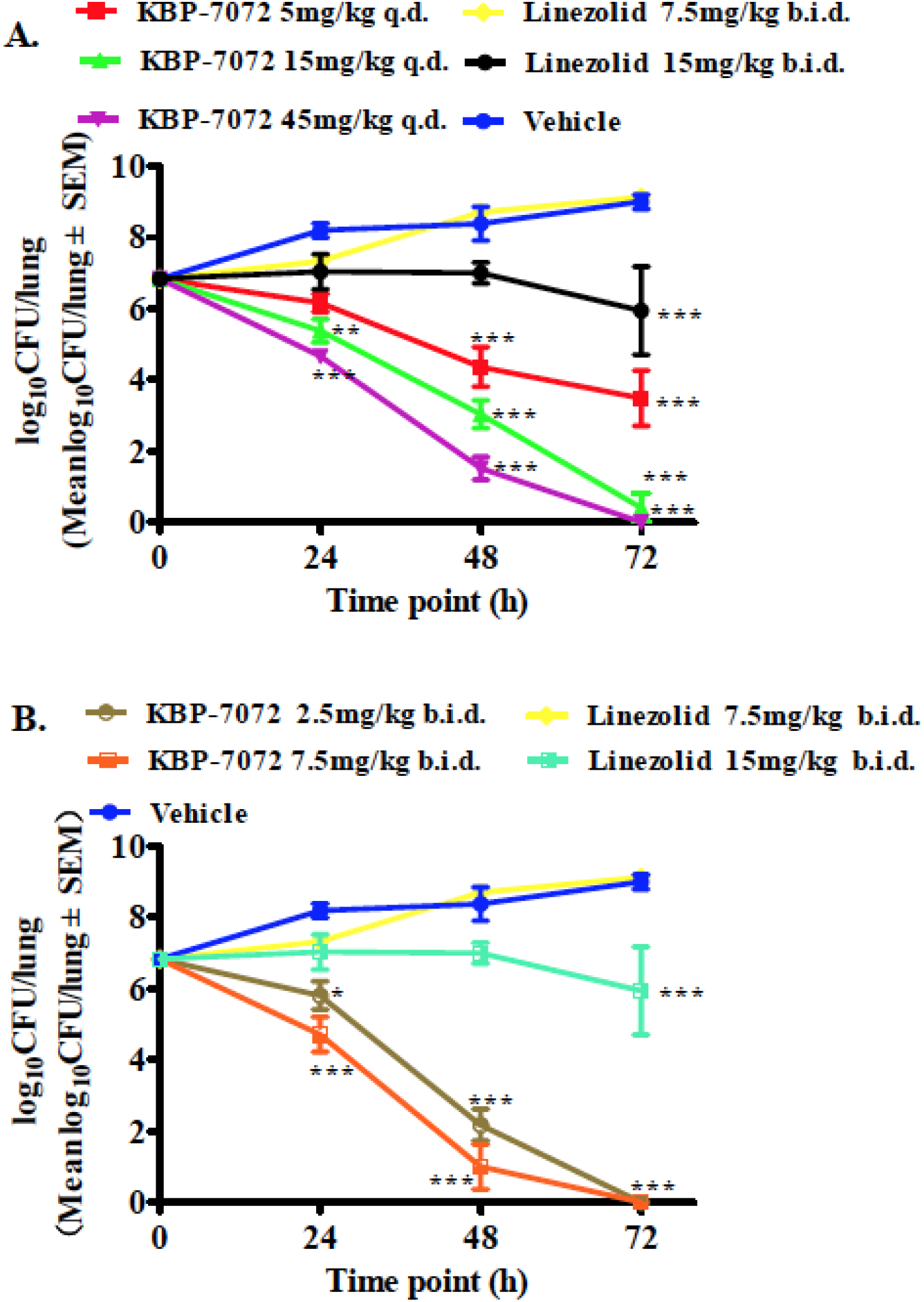
Efficacy of KBP-7072 in mouse pneumonia model infected by intranasal inoculation of *Streptococcus pneumoniae*. A. Colony forming unit in animals treated by KBP-7072 dosed once daily. B. Colony forming unit in animals treated by KBP-7072 dosed twice daily. (* p<0.05; ** p< 0.01; *** p<0.001).

**Figure 4.**
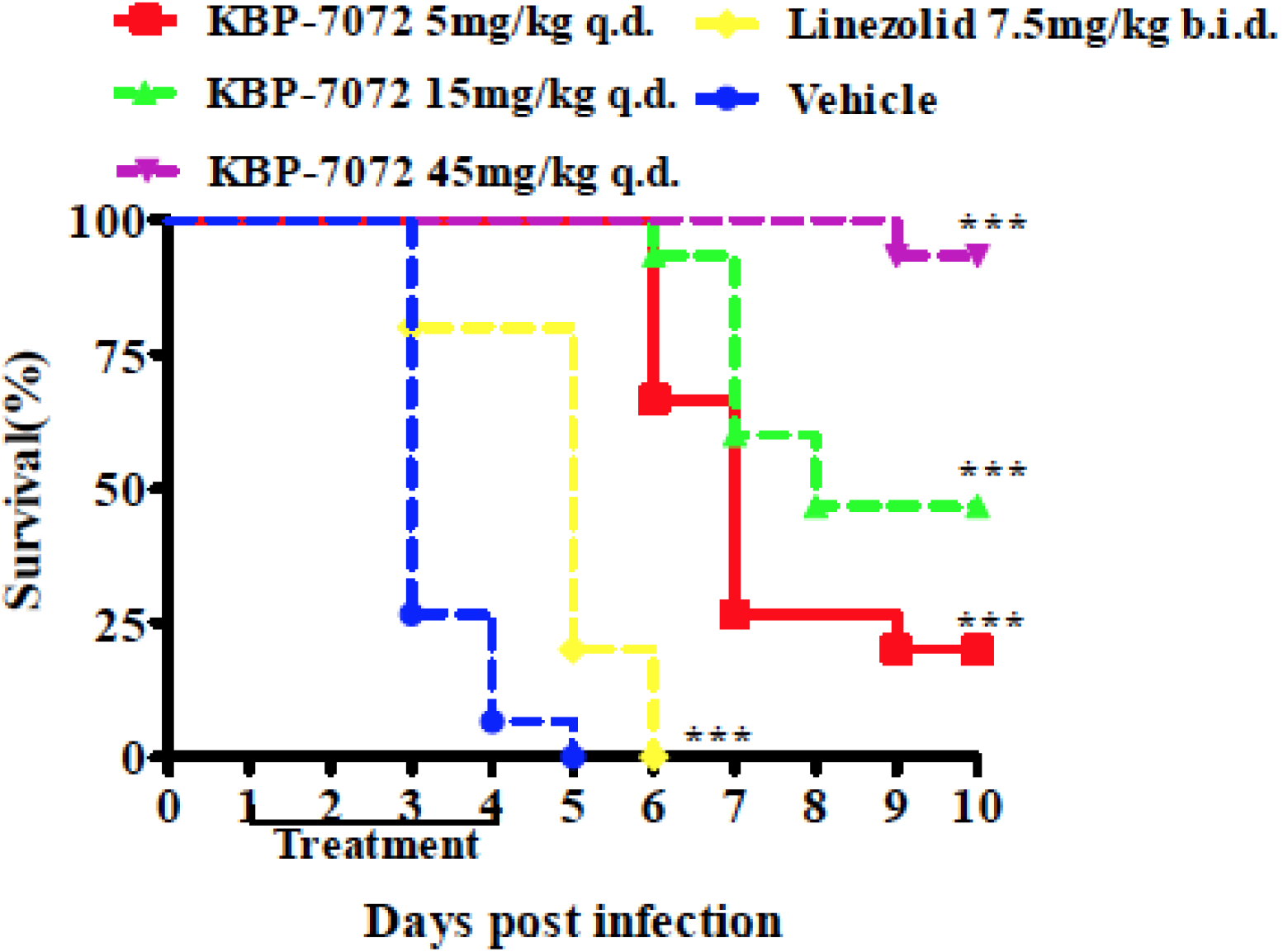
Survival of experimental animals. Log Rank Test was used to analyze survival rate. (* p<0.05; ** p<0.01; *** p<0.001).

### Thigh Infection Model

In the *S. aureus* 6424MRSA-363-induced thigh infection model, KBP-7072 resulted in a dose-dependent reduction of CFUs (**Table 2**). Compared to vehicle, all doses KBP-7072 produced significant (*p*<0.001) reductions of CFUs at 24 hours. In the *K. pneumoniae* 6680kpn-522-induced thigh infection model, KBP-7072 resulted in a dose-dependent reduction in CFUs (**Table 3**). Compared to vehicle, all doses of KBP-7072 resulted in significant (*p*<0.001) reductions in CFUs. The KBP-7072 10 mg/kg bid group showed better activity than the 20 mg/kg qd group against *K. pneumoniae* 6680kpn-522 (*p*<0.001). Similarly, the 20 mg/kg bid group showed better activity than the 40 mg/kg qd group (*p*<0.001). In the *E. coli* 6691eco-558-induced thigh infection model, KBP-7072 resulted in dose-dependent reductions in CFUs (**Table 4**). Compared to vehicle, all doses of KBP-7072 resulted in significant (*p*<0.001) reductions in CFUs. At the once daily dosing schedule, KBP-7072 demonstrated dose dependent reductions in CFUs.

**Table 2.** Efficacy of KBP-7072 in *Staphylococcus Aureus* 6424MRSA-363-Induced Murine Thigh Infection Model.

| Dose<br>(mg/kg) | Schedule | Inoculation<br>Concentration<br>(log<sub>10</sub>CFU/thigh) | 0 h-Control<br>log<sub>10</sub>CFU/thigh<br>Mean ± SD | 24 h<br>log<sub>10</sub>CFU/thigh<br>Mean ± SD | %<br>Reduction * | *p*-<br>value*<br>* |
| --- | --- | --- | --- | --- | --- | --- |
| 10 | qd | 5.79 | 6.41±0.09 | 6.80±0.53 | 25.1 | <0.001 |
|  | bid |  |  | 5.50 ±0.23 | 39.4 | <0.001 |
|  | tid |  |  | 5.62±0.46 | 38.1 | <0.001 |
|  | qid |  |  | 4.88 ±0.34 | 46.3 | <0.001 |
| 20 | qd |  |  | 5.73 ±0.93 | 36.9 | <0.001 |
|  | bid |  |  | 4.75±0.27 | 47.7 | <0.001 |
| 40 | qd |  |  | 5.31±0.26 | 41.6 | <0.001 |
| Vehicle | qd |  |  | 9.08±0.08 | 0 | - |

\* % Reduction = [1-(Log<sub>10</sub>CFU/thing treated ÷ Log<sub>10</sub>CFU/thing vehicle)]×100
\*\* Data analyzed by one-way analysis of variance followed by Tukey test
CFU = colony forming unit; SD = standard deviation

**Table 3.** Activity of KBP-7072 in a *Klebsiella Pneumoniae* 6680kpn-522-Induced Murine Thigh Infection Model.

| Dose<br>(mg/kg) | Schedule | Inoculation<br>Concentration<br>(log<sub>10</sub>CFU/thigh) | 0 h-Control<br>log<sub>10</sub>CFU/thigh<br>Mean ± SD | 24 h<br>log<sub>10</sub>CFU/thigh<br>Mean ± SD | %<br>Reduction* | p-<br>value** |
| --- | --- | --- | --- | --- | --- | --- |
| 10 | qd | 5.76 | 6.60±0.09 | 9.40±0.18 | 9.8 | <0.001 |
|  | bid |  |  | 6.37±0.41 | 38.9 | <0.001 |
| 15 | bid |  |  | 6.04±0.38 | 42.1 | <0.001 |
| 20 | qd |  |  | 8.73±0.16 | 16.3 | <0.001 |
|  | bid |  |  | 5.45±0.12 | 47.7 | <0.001 |
| 30 | qd |  |  | 8.27±0.17 | 20.7 | <0.001 |
| 40 | qd |  |  | 7.51±0.44 | 28.0 | <0.001 |
|  | bid |  |  | 5.04±0.19 | 51.7 | <0.001 |
| Vehicle | qd |  |  | 10.43±0.09 | 0 | - |

\* % Reduction = [1-(Log<sub>10</sub>CFU/thing treated ÷ Log<sub>10</sub>CFU/thing vehicle)]×100
\*\* Data analyzed by one-way analysis of variance followed by Tukey test
CFU = colony forming unit; SD = standard deviation

**Table 4.** Efficacy of KBP-7072 in *Escherichia Coli* 6691eco-558-induced Murine Thigh Infection Model.

| Dose<br>(mg/kg) | Schedule | Inoculation<br>Concentration<br>(log<sub>10</sub>CFU/thigh) | 0 h-Control<br>log<sub>10</sub>CFU/thigh<br>Mean ± SD | 24 h<br>log<sub>10</sub>CFU/thigh<br>Mean ± SD | %<br>Reduction* | *p*<br>Value** |
| --- | --- | --- | --- | --- | --- | --- |
| 15 | qd | 6.06 | 6.67±0.03 | 8.45±0.08 | 15.9 | <0.001 |
|  | tid |  |  | 5.31±0.40 | 47.2 | <0.001 |
| 20 | qid |  |  | 5.00±0.24 | 50.3 | <0.001 |
| 30 | qd |  |  | 8.08±0.24 | 19.6 | <0.001 |
|  | bid |  |  | 5.17±0.11 | 48.5 | <0.001 |
|  | qid |  |  | 4.98±0.13 | 50.5 | <0.001 |
| 40 | qd |  |  | 7.66±0.24 | 23.7 | <0.001 |
|  | tid |  |  | 5.06±0.24 | 49.6 | <0.001 |
| 50 | qd |  |  | 7.00±0.26 | 30.4 | <0.001 |
| Vehicle | qd |  |  | 10.05±0.16 | 0 | - |

\* % Reduction= [1-(Log<sub>10</sub>CFU/thing treated ÷ Log<sub>10</sub>CFU/thing vehicle)]×100
\*\*Data analyzed by one-way analysis of variance followed by Tukey test
CFU = colony forming unit; SD = standard deviation

### Relationship Between Efficacy and PK/PD Parameters of KBP-7072

In the KBP-7072-treated thigh infection models, the relationship between the log_10_ CFU change and the PK/PD parameters (the ratio of the area under the unbound concentration – time curve to the minimum inhibitory concentration [24-h *f*AUC/MIC], the ratio of the unbound peak plasma concentration to the MIC [*f*C_max_/MIC], and the percentage of the dosing interval that the unbound drug concentration exceeded the MIC [%T>MIC_free_]) was examined. The *f*AUC/MIC ratio was the most predictive PK/PD parameter for the efficacy of KBP-7072 *in vivo* (**Figures 5-7 and Tables 6-8**). The magnitude of the PK/PD parameters associated with each dose was calculated from the following equation: E=E_0_−E_max_×(PK parameter)N/(EC_50_^N^+(PK parameter)^N^). The relationship between KBP-7072 MIC and *f*AUC/MIC necessary to achieve a static effect, 1 log_10_ killing, and 2 log_10_ killing is shown in **Table 5**. As a measure of the *in vivo* efficacy of KBP-7072 against *S. aureus, K. pneumoniae*, and *E. coli* clinical isolates, the mean *f*AUC/MIC ratio was 4.39 for achieving a bacteriostatic effect, and 8.34 and 13.46 for achieving a bactericidal effect.

**Figure 5.**
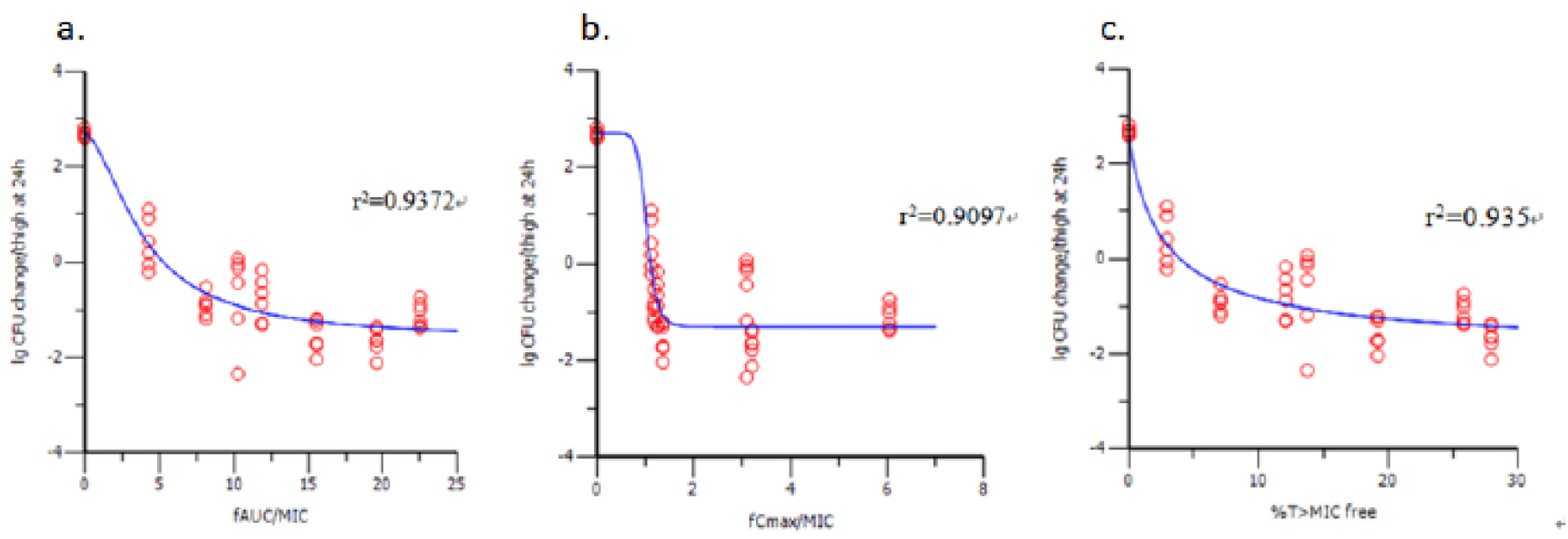
Relationship between the log_10_ CFU change and the PK/PD parameters 24-h *f*AUC/MIC (a), *f*Cmax/MIC (b) and %T>MIC_free_ (c) in KBP-7072-treated methicillin-resistant *Staphylococcus aureus* 6424MRSA-363-induced thigh infection model. (*f*AUC/MIC Parameter: Emax=4.2851363, EC50=3.9220294, E0=2.6738137, N=1.7180344).

**Figure 6.**
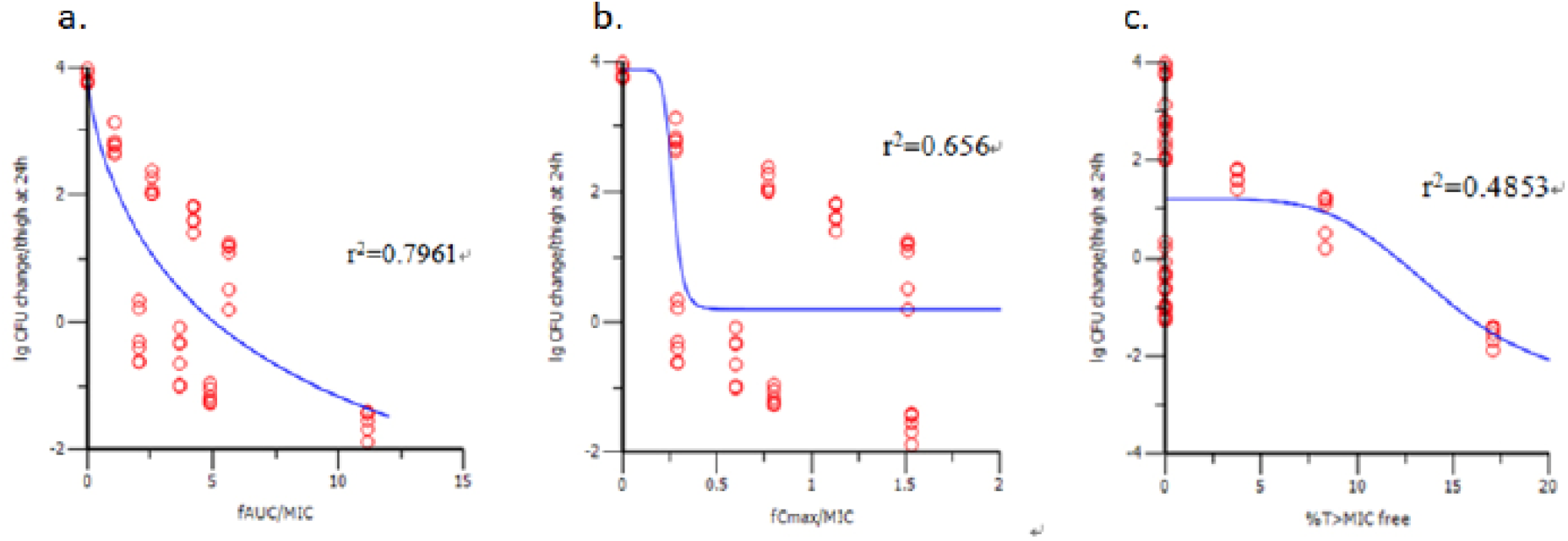
Relationship between the log10 CFU change and the PK/PD parameters 24-h *f*AUC/MIC (a), *f*Cmax/MIC (b) and %T>MIC_free_ (c) in KBP-7072-treated *Klebsiella pneumoniae* 6680kpn-522-induced thigh infection model. (*f*AUC/MIC Parameter: Emax=9.4409802, EC50=8.2909711, E0=3.8799326, N=0.72923001).

**Figure 7.**
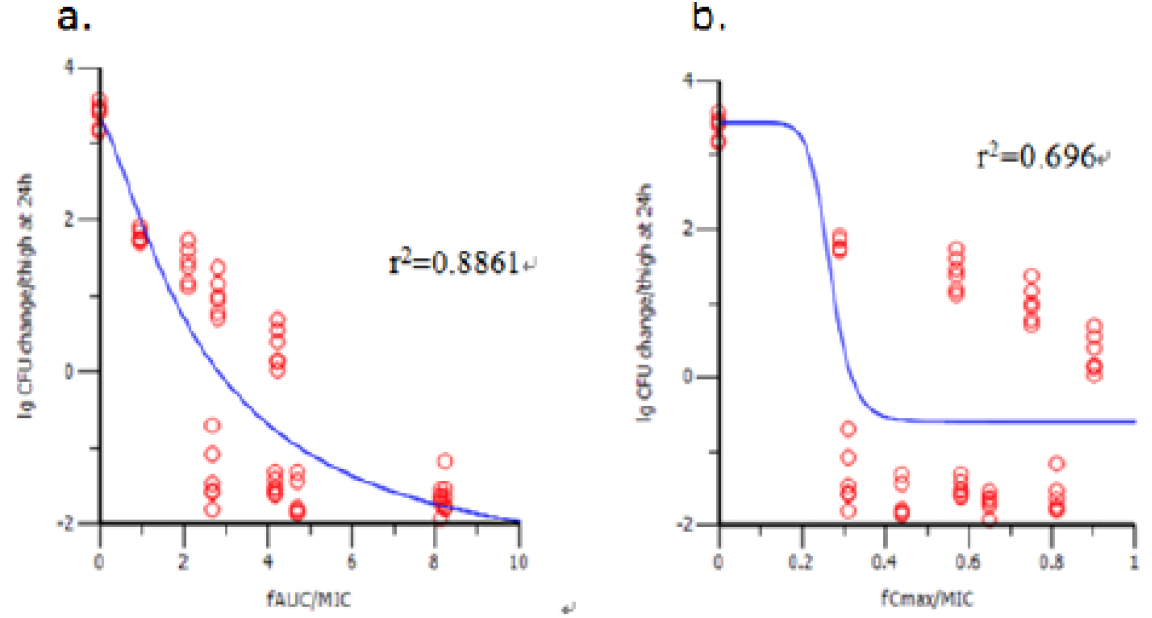
Relationship between the log10 CFU change and the PK/PD parameters 24-h *f*AUC/MIC (a) and *f*Cmax/MIC (b) in KBP-7072-treated *Escherichia coli* 6691eco-558-induced thigh infection model. (*f*AUC/MIC Parameter: Emax=6.0756719, EC50=2.4501195, E0=3.3397329, N=1.3692287).

**Table 5.** Relationship Between KBP-7072 MIC and *f*AUC/MIC Necessary for Achieving a Static Effect and 1 log_10_ and 2 log_10_ Killing.

| Isolate | MIC (µg/mL) | *f*AUC/MIC |  |  |
| --- | --- | --- | --- | --- |
|  |  | static effect | 1 log<sub>10</sub>kill | 2 log<sub>10</sub>kill |
| 6424MRSA-363 | 0.25 | 5.27 | 11.14 | NA* |
| 6680 *K. pneumoniae*-522 | 1 | 5.06 | 9.10 | 16.49 |
| 6691 *E. coli*-558 | 2 | 2.83 | 4.78 | 10.42 |
| Mean | / | 4.39 | 8.34 | 13.46 |

\*NA: 2 log<sub>10</sub>kill was achieved.

**Table 6.** PK/PD Parameters* of KBP-7072 in *Staphylococcus aureus* (No: 6424MRSA-363)-induced Murine Thigh Infection Model.

| Dose (mg/kg) | Interval | $fAUC/MIC^*$ | $fC_{max}/MIC^*$ | $\%T > MIC_{free}^*$ |
| --- | --- | --- | --- | --- |
| 10 | qd | 4.28 | 1.12 | 2.92 |
|  | bid | 8.15 | 1.17 | 7.08 |
|  | tid | 11.89 | 1.25 | 12.08 |
|  | qid | 15.55 | 1.36 | 19.17 |
| 20 | qd | 10.26 | 3.09 | 13.75 |
|  | bid | 19.58 | 3.20 | 27.92 |
| 40 | qd | 22.50 | 6.04 | 25.83 |

\* PK/PD parameters were calculated using WinNonlin Phoenix 6.1.0 software with the MIC value, PK, and plasma protein binding data.

**Table 7.** PK/PD Parameters* of KBP-7072 in *Klebsiella pneumoniae* (No: 6680kpn-522) in a Murine Thigh Infection Model.

| Dose (mg/kg) | Interval | $fAUC/MIC^a$ | $fC_{max}/MIC^a$ | %T>MIC<sub>free</sub><sup>a</sup> |
| --- | --- | --- | --- | --- |
| 10 | qd | 1.07 | 0.28 | 0.00 |
|  | bid | 2.04 | 0.29 | 0.00 |
| 15<sup>b</sup> | bid | 3.67 | 0.60 | 0.00 |
| 20 | qd | 2.56 | 0.77 | 0.00 |
|  | bid | 4.89 | 0.80 | 0.00 |
| 30<sup>c</sup> | qd | 4.22 | 1.13 | 3.75 |
| 40 | qd | 5.62 | 1.51 | 8.33 |
|  | bid | 11.15 | 1.53 | 17.08 |

<sup>a</sup> PK/PD parameters were calculated using WinNonlin Phoenix 6.1.0 software with the MIC value, PK, and plasma protein binding data.
<sup>b, c</sup> The PK parameter of KBP-7072 after single dose subcutaneous administration was linear.
The PK/PD parameter for dose 15 mg/kg and 30 mg/kg was calculated using the PK data of 20 mg/kg and 40 mg/kg, respectively.

**Table 8.**
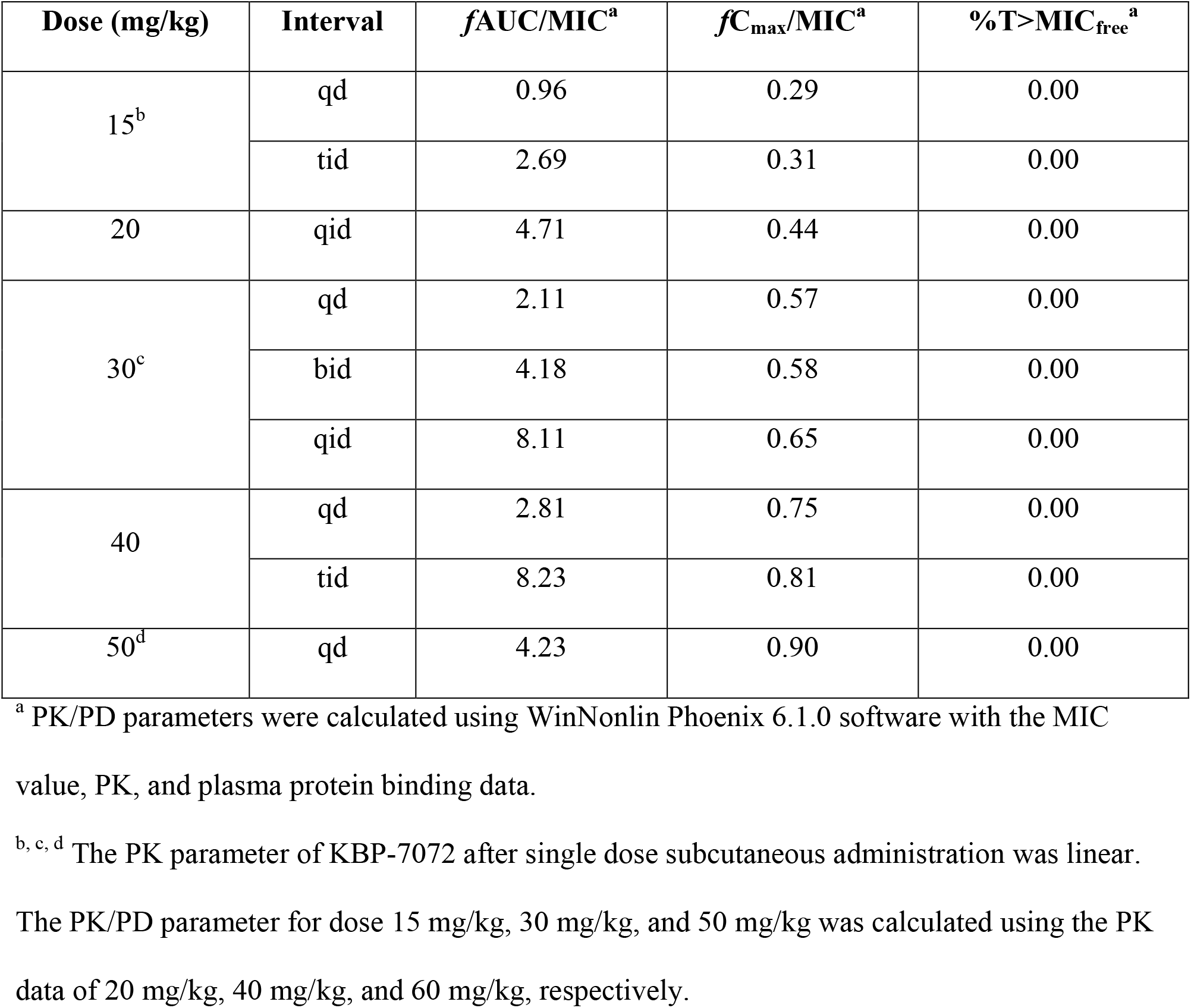
PK/PD Parameters of KBP-7072 Against *Escherichia coli* (No: 6691eco-558) in a Murine Thigh Infection Model.

| Dose (mg/kg) | Interval | $fAUC/MIC^a$ | $fC_{max}/MIC^a$ | %T>MIC<sub>free</sub><sup>a</sup> |
| --- | --- | --- | --- | --- |
| 15<sup>b</sup> | qd | 0.96 | 0.29 | 0.00 |
|  | tid | 2.69 | 0.31 | 0.00 |
| 20 | qid | 4.71 | 0.44 | 0.00 |
| 30<sup>c</sup> | qd | 2.11 | 0.57 | 0.00 |
|  | bid | 4.18 | 0.58 | 0.00 |
|  | qid | 8.11 | 0.65 | 0.00 |
| 40 | qd | 2.81 | 0.75 | 0.00 |
|  | tid | 8.23 | 0.81 | 0.00 |
| 50<sup>d</sup> | qd | 4.23 | 0.90 | 0.00 |

<sup>a</sup> PK/PD parameters were calculated using WinNonlin Phoenix 6.1.0 software with the MIC value, PK, and plasma protein binding data.
<sup>b, c, d</sup> The PK parameter of KBP-7072 after single dose subcutaneous administration was linear.
The PK/PD parameter for dose 15 mg/kg, 30 mg/kg, and 50 mg/kg was calculated using the PK data of 20 mg/kg, 40 mg/kg, and 60 mg/kg, respectively.

## DISCUSSION

Results from the studies reported here demonstrated significant reductions in bacterial growth in pneumonia infection models. KBP-7072 had greater dose-dependent antibacterial activity compared to minocycline and significantly increased the survival rate and prolonged median survival of treated animals in a *K. pneumoniae*-induced murine pneumonia model. Similar results were observed with KBP-7072 compared to linezolid in a *S. pneumoniae*-induced murine pneumonia model. The activity and PK/PD relationship of KBP-7072 in a murine thigh infection model revealed that KBP-7072 resulted in dose-dependent reductions of CFUs, with higher total doses of KBP-7072 resulting in greater reductions of CFUs. In a MRSA-induced thigh infection model using the same total dose, more frequent dosing of KBP-7072 did not achieve higher efficacy than once daily dosing, which supports a once daily dosing as the clinical dosing regimen. In the murine thigh model, the antibacterial activity of KBP-7072 was correlated with the time-dependent PK/PD parameters, *f*AUC/MIC and %T>MIC_free_, although the *f*AUC/MIC ratio was the most predictive PK/PD parameter for KBP-7072 for demonstrating *in vivo* activity. The mean *f*AUC/MIC ratio required for achieving a bacteriostatic effect of KBP-7072 was 4.39 and for achieving 1-log_10_ kill and 2-log_10_ kill was 8.34 and 13.46, respectively. Previous *in vitro* studies with KBP-7072 demonstrated MIC_90_ values <1 μg/mL across a range of Gram-negative and Gram-positive pathogens, including multidrug resistant and typical and atypical pathogens associated with CABP (Huband et al, 2020).15

The efficacy of different classes of antibiotics is most often predicted by time- or dose-dependent PK/PD parameters. For tetracyclines, time-dependent parameters, i.e., AUC/MIC, are most often predictive of antimicrobial bacteriostatic and bactericidal activity.^16^ Results from *in vivo* infection models with other antibiotics of the tetracycline class demonstrate that the time plasma concentrations of drug are above the MIC or AUC/MIC is the optimal PK/PD parameter for establishing efficacy.^17–23^ In a PK/PD evaluation of KBP-7072 in a murine model of pneumonia, plasma AUC/MIC values for a 2-log_10_ kill were 7.2 and 31.4 for *Staphylococcus aureus* and *Streptococcus pneumoniae* at 24 hours.^24^ Peak KBP-7072 concentrations ranged from 0.12 to 25.2 mcg/mL.^24^ Epithelial lining fluid (ELF) concentrations with KBP-7072 were 82% to 238% of plasma concentrations. While blood and ELF levels were not obtained in the pneumonia models reported here, results from previous studies in thigh infection models allowed calculation of PK/PD parameters for KBP-7072, and results were comparable with those previously reported.^24^

Limitations of these results include that no blood or epithelial lining fluid concentrations were obtained directly from animals studied. Consequently, PK/PD calculations were based on previous published work with KBP-7072. Nevertheless, prior PK results and results from in vivo testing reported here were obtained in murine pneumonia models using CD-1 mice. Thus, these results provide preliminary evidence of an effective dose of KBP-7072 for treating pneumonia and provide support for further nonclinical and clinical studies to determine the optimal dose of KBP-7072 for serious infections.

KBP-7072 is undergoing clinical development for treating CABP and other serious infections due to Gram-positive and Gram-negative aerobes including many multidrug resistant pathogens. Results from these studies of two different *in vivo* models of infection across multiple pathogens support once daily administration of KBP-7072 and are consistent with findings from single and multiple dose studies of KBP-7072 in healthy volunteers where the elimination half-life exceeded 24 hours.^13,14^ These results together with results from *in vitro* studies of microbiological activity suggest that KBP-7072 is a promising antibiotic with the potential to expand the armamentarium of drugs available to treat serious infections, especially in an era of growing bacterial resistance to antimicrobials.

## METHODS

All *in vivo* studies were conducted under appropriate Institutional Animal Care and Use Committee-approved protocols and in accordance with under KBP BioSciences Institutional Animal Care and Use Committee guidelines. Animals were 6-8-week-old female CD-1 mice, 23 - 27 g obtained from Beijing Vital River Laboratory Animal Technology Co. Limited and housed in a segregated pathogen-free room under controlled temperature, humidity, airflow, and lighting conditions.

### Pneumonia Infection Models

For the pneumonia infection model, animals were randomized to treatment groups (15 per group) based on body weight. After randomization, animals were rendered neutropenic with intraperitoneal (ip) cyclophosphamide 150 mg/kg at Day −4 and Day −1, then anesthetized with 6.5% pentobarbital sodium 65 mg/kg ip. Each animal, under light anesthesia, was inoculated with 50 μL of a log phase culture of *K. pneumoniae* 5615kpn-493 or *S. pneumoniae* 6962spn-310 placed on the tip of the nares. The second subculture of *S. pneumoniae* 6962spn-310 strains was prepared less than 20 hours before inoculation. Prior to inoculation, a suspension of 10^8^ CFU of *S. pneumoniae* per mL was prepared in Mueller-Hinton (MH) broth (including 10% bovine serum) by adjusting to a 2.5 McFarland turbidity standard. Final inoculum densities (CFU per milliliter) were confirmed by serial dilution and culture of each inoculum.

For the *K. pneumoniae* model, *in vitro* activity (MIC) against a minocycline- and tetracycline-resistant strain of *K. pneumoniae* was 1, 4, and 8 μg/mL for KBP-7072, minocycline, and tetracycline, respectively, using CLSI standards.^25,26^ For the *S. pneumoniae* model, *in vitro* activity (MIC) against a minocycline- and tetracycline-resistant strain of *S. pneumoniae* was 0.03, 1, 8, and >16 μg/mL for KBP-7072, linezolid, minocycline, and tetracycline, respectively.

For the *K. pneumoniae* model, animals were dosed with oral KBP-7072 150, 300, and 600 mg/kg qd or 75 or 150 mg/kg bid; oral minocycline 300 mg/kg qd; or oral vehicle 0.2 mL qd. KBP-7072 and comparator antibiotics were started 3 hours post inoculation and continued for 3 days. For the *S. pneumoniae* model, animals received oral KBP-7072 at doses of 5, 15, or 45 mg/kg, qd or 2.5 or 7.5 mg/kg, bid; oral linezolid 7.5 or 15 mg/kg, bid; and oral vehicle 0.2 mL, qd. Fifteen mice were assigned to each dose group. KBP-7072 and comparator antibiotics were started 18 hours post inoculation and continued for 3 days. Treatment groups were fasted 12 hours before the initiation of dosing. Food was provided 1 hour after the initiation of the dosing. Mice in the bid treatment groups were also fasted 8 hours before the second dosing, and the feeding was resumed 1 hour after dosing. Mice had free access to water throughout the study.

One group of animals was euthanized at 0 (control group), 24, 48 or 72 hours post final dose. The lungs from 6 mice in each group were harvested, homogenized in saline, and serial dilutions of the homogenates were cultured overnight on a MH agar plate. Bacterial CFUs)were presented as log_10_CFU/lung. For a second group of animals, cumulative survival rate after 72 hours of therapy was assessed for 10 days post-infection.

Data on the bacterial colony count of the lungs was analyzed using one-way analysis of variance (ANOVA) followed by Tukey’s Multiple Comparison Test. The survival rate was analyzed by Log Rank Test with comparisons to the vehicle group at same time point.

### Thigh Infection Model

KBP-7072 *in vivo* activity was tested on three clinical bacteria isolates collected from Chinese hospitals in a murine thigh infection model: one MRSA isolate (6424MRSA-363), one *K. pneumoniae* isolate (6680kpn-522), and one *E. coli* isolate (6691eco-558). The *in vitro* activity of KBP-7072 vs. minocycline was determined for each of the 3 isolates using CLSI standards.^25,26^

To compromise the immune system, mice were injected ip with cyclophosphamide 150 mg/kg (10 mL/kg of 15 mg/mL stock solution) on Day 4 before bacteria inoculation and 100 mg/kg (10 mL/kg of 10 mg/mL stock solution) on Day 1 before bacteria inoculation. Two hours prior to the initiation of antimicrobial therapy, each thigh of the neutropenic mouse was inoculated intramuscularly with a 0.1 mL solution containing approximately 10^6^ CFU/thigh of the test isolate prepared from a fresh subculture.

Two hours post infection; mice were treated subcutaneously based on the body weight with KBP-7072 or vehicle. For *S. aureus*, mice (n=3/group) were randomized to vehicle or KBP-7072 10 mg/kg qd, bid, tid or qid; 20 mg/kg qd or bid; or 40 mg/kg qd. For *K. pneumoniae*, mice received vehicle or KBP-7072 doses of 10 mg/kg qd or bid; 15 mg/kg bid; 20 mg/kg qd or bid; 30 mg/kg qd; or 40 mg/kg qd or bid. For *E. coli*, mice received vehicle or KBP-7072 doses of 15 mg/kg qd or tid; 20 mg/kg qid; 30 mg/kg qd, bid or qid; 40 mg/kg qd or tid; or 50 mg/kg qd.

At 24 hours post initial dose, mice were euthanized by CO_2_ exposure and followed by cervical dislocation. Thighs were cleaned with 70% ethanol, skins were removed and infected muscles from knee to hip joint were harvested under sterile condition. Each muscle was mixed with 5 mL saline and was homogenized at 20,000 rpm. The homogenates were serially diluted, and 50 μL of three different serial diluted homogenates of each thigh were placed on MH agar plates at 35°C for 18 hours for CFU determination. One group of three mice was harvested at 0 hour to serve as the baseline control group.

Activity, defined as the change in bacterial density, was calculated as the log_10_ change of bacterial CFU per thigh: Log_10_ change of CFU per thigh = Log_10_ CFU of treated group-log_10_ CFU of the baseline control group. To evaluate the PK/PD relationship of KBP-7072 in the *in vivo* disease model, data were used from a study of KBP-7072 conducted in cyclophosphamide-induced neutropenic CD-1 mice after a single subcutaneous injection of KBP-7072 doses of 60, 40, 20, 10, 5, and 2.5 mg/kg, respectively to establish PK parameters.^27^

Efficacy (log_10_CFU change/thigh) was measured by the arithmetic mean change in log_10_CFU per thigh of the 24 hours control or treatment groups from the 0-h baseline control mouse (2 hours after inoculation). PK/PD parameters *f*AUC/MIC, *f*C_max_/MIC, and %T>MIC_free_ were calculated by the WinNonlin Phoenix 6.1.0 software, using PK data, plasma protein binding, and MIC values.^27^ The diagram of efficacy (log_10_CFU change/thigh as ordinate) and PK/PD parameters (*f*AUC/MIC, *f*C_max_/MIC, and %T>MIC_free_ as abscissa) was plotted. The magnitude of the PK/PD parameters associated with each dose was calculated from the following equation: E=E_0_-E_max_×(PK parameter)^N^/(EC_50_^N^+(PK parameter)^N^),^28^ and PK/PD parameters were calculated when efficacy was static effect, 1 and 2 log_10_ reductions in colony counts compared to the numbers at the start of therapy.

## ACKNOWLEDGEMENT

The authors would like to acknowledge the editorial assistance of Richard S. Perry, PharmD in the development of this manuscript, which was supported by KBP Biosciences, USA Inc., Princeton, NJ.

## Author Contributions

All authors contributed to data analysis and interpretation, reviewed the manuscript for intellectual content, and approval submission of the manuscript.

